# Loss of Glucose Transporter 1 in Mouse Uterine Cells Disrupts Trophoblast Differentiation and Promotes Gestational Diabetes

**DOI:** 10.64898/2026.06.04.730181

**Authors:** Ritwik Shukla, Athilakshmi Kannan, Kevin W. Porter, Cheyenne S. Summers, Arpita Bhurke, Milan K. Bagchi, Indrani C. Bagchi

## Abstract

A successful pregnancy hinges on a finely coordinated dialogue between the maternal endometrium and the implanting embryo. Following embryo attachment to the uterine epithelium, underlying stromal cells undergo a transformation into decidual cells that promote a vascularized maternal-fetal interface and direct trophoblast lineage decisions through paracrine cues. However, the metabolic adaptations that enable decidual cells to support these energetically demanding processes remain poorly understood. Here, using a uterine-specific knockout mouse model, we identify Glucose Transporter 1 (Glut1) as a critical metabolic regulator linking endometrial glucose uptake to reproductive success. We demonstrate that stromal Glut1, induced by hypoxia-inducible factor 2α (Hif2α), sustains a Hif2α-Rab27b feed-forward circuit that drives vesicular trafficking during pregnancy through the glucose-sensing transcription factor MAX-like protein X (Mlx). Mice lacking endometrial Glut1 are severely subfertile despite normal embryo attachment. Glut1-deficient uteri exhibit impaired stromal extracellular vesicle secretion, defective decidual angiogenesis, and marked dysregulation of trophoblast differentiation, including expansion of trophoblast progenitors, accumulation of glycogen trophoblast cells, and altered placental lactogen production. These placental defects culminate in mid-gestation fetal loss and maternal gestational diabetes mellitus (GDM). Collectively, our findings establish endometrial Glut1 as a metabolic gatekeeper of maternal glucose homeostasis and placentation and introduce a genetically tractable mouse model of spontaneously developing GDM, a disorder affecting nearly one in seven pregnancies worldwide.

**SIGNIFICANCE:** This study uncovers endometrial Glut1 as a previously unrecognized metabolic regulator of placentation, demonstrating that maternal stromal glucose uptake dictates trophoblast fate and maternal glycemic control, and providing the field with its first genetically defined mouse model of spontaneously arising gestational diabetes.

## INTRODUCTION

A successful pregnancy is among the most metabolically demanding processes a mammalian organism undertakes. From the moment the embryo attaches to the uterine luminal epithelium and embeds into the underlying stroma, the maternal endometrium must mount an extraordinary set of cellular and metabolic adaptations to ensure a continuous supply of nutrients and signaling cues to the developing conceptus (1, 38). Central to these adaptations is decidualization, the proliferation and morphological transformation of endometrial stromal cells into specialized decidual cells, accompanied by the rapid expansion of an angiogenic vascular network that supports the growing embryo (2–4, 39). In addition, the decidua transmits maternal cues to the developing trophoblast, thereby shaping placental architecture and function (5, 6, 40). When this exquisitely tuned program goes awry, the consequences are profound, ranging from implantation failure and recurrent pregnancy loss to intrauterine growth restriction, preeclampsia, and gestational diabetes mellitus (7, 8, 41, 42). A central, unresolved question in reproductive biology is how decidual cells reprogram their metabolism, particularly their handling of glucose, the dominant fuel of the periimplantation uterus to power these critical events.

In a previous study, we reported that Hif2α (hypoxia-inducible factor 2 alpha) induced at the time of embryo implantation plays a critical role in establishing pregnancy (9). Hif2α enables cells to sense and adapt to hypoxic microenvironments by reprogramming metabolism, including upregulation of glucose transporters and glycolytic enzymes (10). Comparative gene expression profiling of *Hif2α*-intact and *Hif2α*-null endometrial stromal cells identified Glucose Transporter 1 (Glut1) as a key downstream target induced during early pregnancy. The Gluts (encoded by *Slc2a* genes) constitute a family of facilitative membrane transporters comprising 14 members in mammals (11, 12), of which Glut1 is the only family member regulated by Hif2α in the uterus during implantation. Notably, downregulation of GLUT1 in the human endometrium has been clinically associated with idiopathic infertility (13), yet how endometrial GLUT1 contributes to pregnancy at the cellular and molecular levels remains largely unknown.

In the present study, we use a conditional genetic approach to interrogate the function of endometrial Glut1 in pregnancy. We show that uterine-specific ablation of *Glut1* produces a striking subfertility phenotype. Although embryos attach normally and stromal cells initiate decidualization, late-stage stromal differentiation, decidual angiogenesis, and stromal extracellular vesicle secretion are all severely compromised in the absence of Glut1. These maternal defects, in turn, derail trophoblast differentiation and placental architecture, leading to mid-gestation fetal loss, intrauterine growth restriction, and a sustained elevation of maternal blood glucose from mid-gestation through term, hallmarks of gestational diabetes. Together, these findings establish endometrial Glut1 as a central metabolic regulator that coordinates decidual function, placentation, and maternal glucose homeostasis during pregnancy, and they provide a clinically informative mouse model of spontaneous pregnancy-specific diabetes mellitus caused by a maternal genetic perturbation.

## RESULTS

### Glut1 is induced downstream of Hif2α signaling in mouse endometrial stromal cells during decidualization

Our previous studies revealed that Hif2α regulates embryo implantation and the establishment of pregnancy (9). To dissect the molecular underpinnings of this control, we isolated RNA from *Hif2α*-intact (*Hif2α^f/f^*) and *Hif2α*-null (*Hif2α^d/d^*) stromal cells and performed gene expression profiling (9). These analyses revealed that the expression of Glucose Transporter 1 (Glut1) was markedly downregulated in *Hif2α*-null stromal cells. Consistent with these data, we observed robust GLUT1 protein expression in stromal cells surrounding the implanted embryo of *Hif2α^f/f^* uteri on day 5 of gestation (Fig. 1, left), whereas its expression was dramatically reduced in *Hif2α^d/d^* stromal cells (Fig. 1, right).

**Figure 1.**
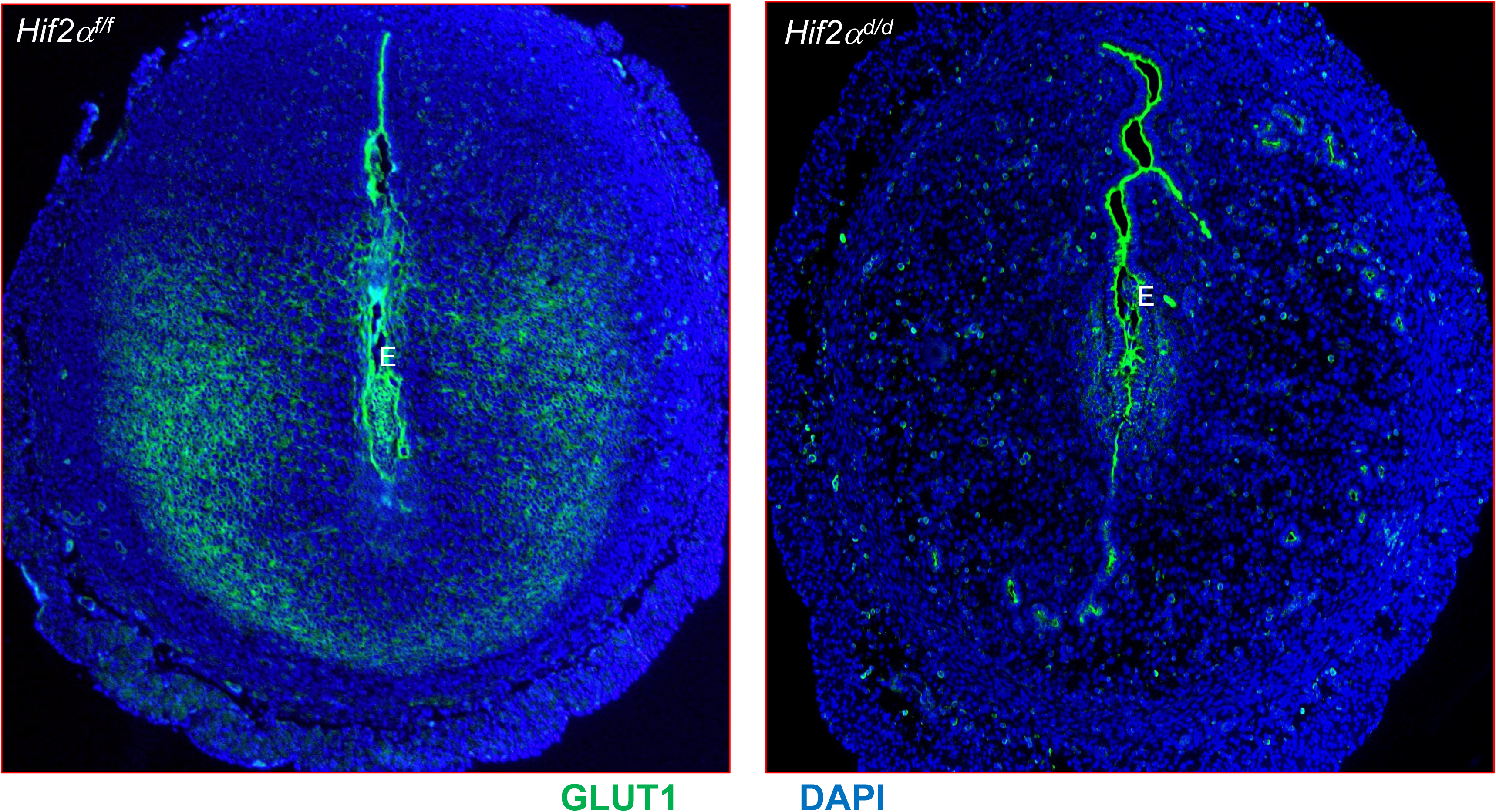
Hif2α induces Glut1 expression in uterine stromal cells during implantation. IF analysis of GLUT1 (green) in *Hif2α^f/f^* and *Hif2α^d/d^* uteri on day 5 of pregnancy. N=3 mice were examined. E denotes an embryo.

We next examined the spatial distribution of GLUT1 across days 5 to 8 of pregnancy (Fig. 2, panels a-f). GLUT1 was detected in sub-luminal stromal cells at the implantation site on day 5 (panel a), as well as in the endometrial glands and the implanting embryo. As pregnancy progressed from day 5 to day 8, GLUT1 expression expanded into the secondary decidual zone (panels b-f) and into the mesometrial stroma, where the angiogenic networks supporting early pregnancy develop (panels d-f). These results demonstrate that GLUT1 is progressively expressed in differentiating stromal cells during early pregnancy.

**Figure 2.**
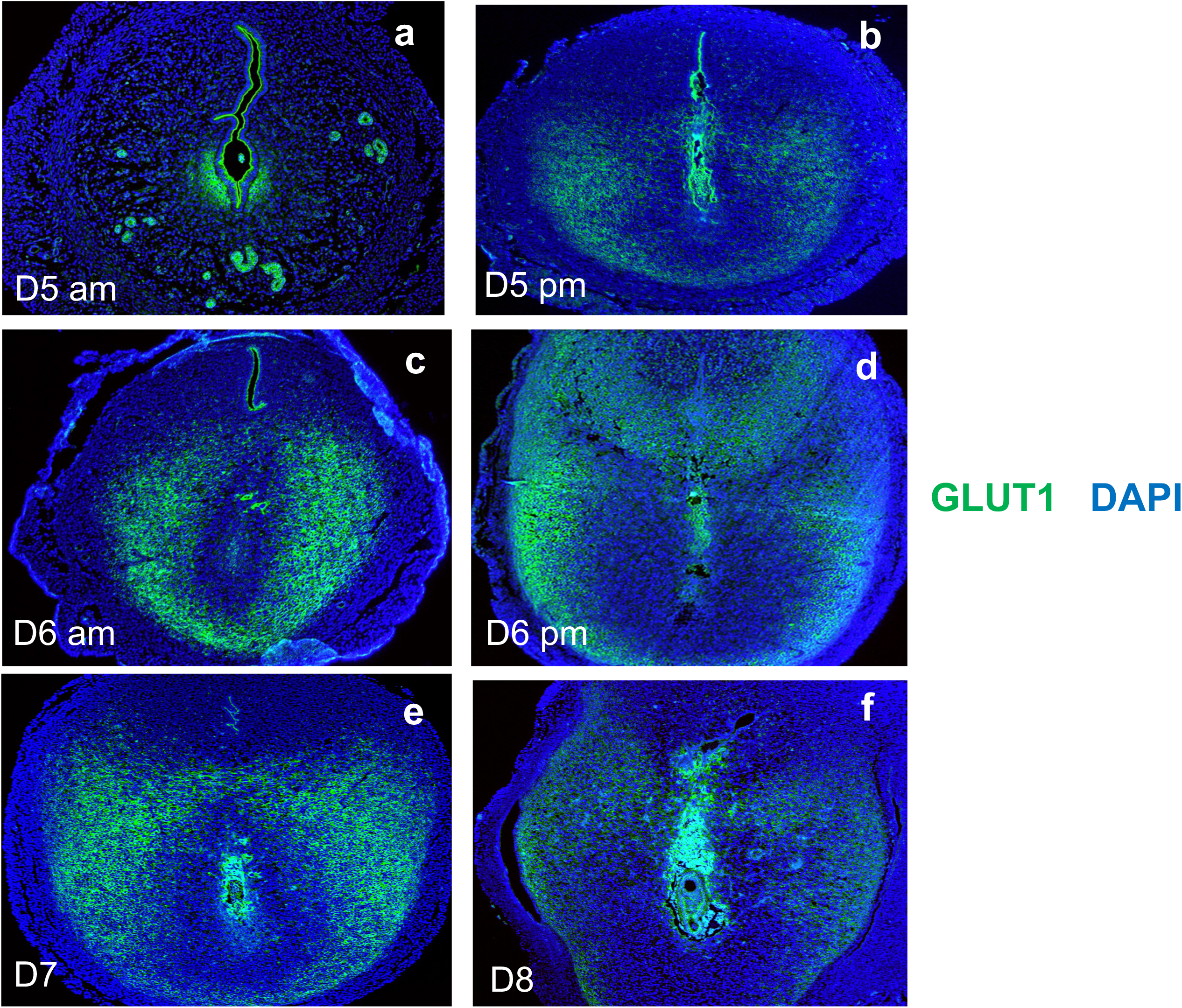
Expression of GLUT1 during early pregnancy overlaps with the decidual phase of gestation. Uterine sections on days 5 morning (a), 5 evening (b), 6 morning (c), 6 evening (d), 7 (e), and 8 (f) of pregnancy were subjected to immunohistochemical analysis using an anti-GLUT1 antibody. E indicates an embryo. Blue: DAPI. Data were repeated with N=2 mice. Representative images are shown.

### Conditional deletion of Glut1 in the uterus leads to severe infertility

To interrogate the function of Glut1 in the uterus, we conditionally deleted the *Glut1* gene in adult mice by crossing mice harboring a “floxed” *Glut1* allele (*Glut1^f/f^*) with *PgrCre/+* mice to generate *Glut1^d/d^* animals (14). The extent of Glut1 deletion was assessed by qPCR and immunofluorescence (IF) (Fig. 3). *Glut1* transcripts were nearly abolished in day 7 pregnant *Glut1^d/d^* uteri, confirming efficient gene ablation (Fig. 3, left). Consistent with the RNA profile, GLUT1 protein was prominent in uterine stromal cells of control *Glut1^f/f^* mice on day 7 of pregnancy but was dramatically reduced in stromal cells of *Glut1^d/d^* uteri (Fig. 3, middle). Because *Pgr-Cre* is not expressed in germ cells, the “floxed” *Glut1* allele remains intact in embryos derived from *Pgr-Cre* dams crossed with wild-type males. Accordingly, GLUT1 expression was retained in implanting embryos within *Glut1^d/d^* uteri (Fig. 3, middle).

**Figure 3.**
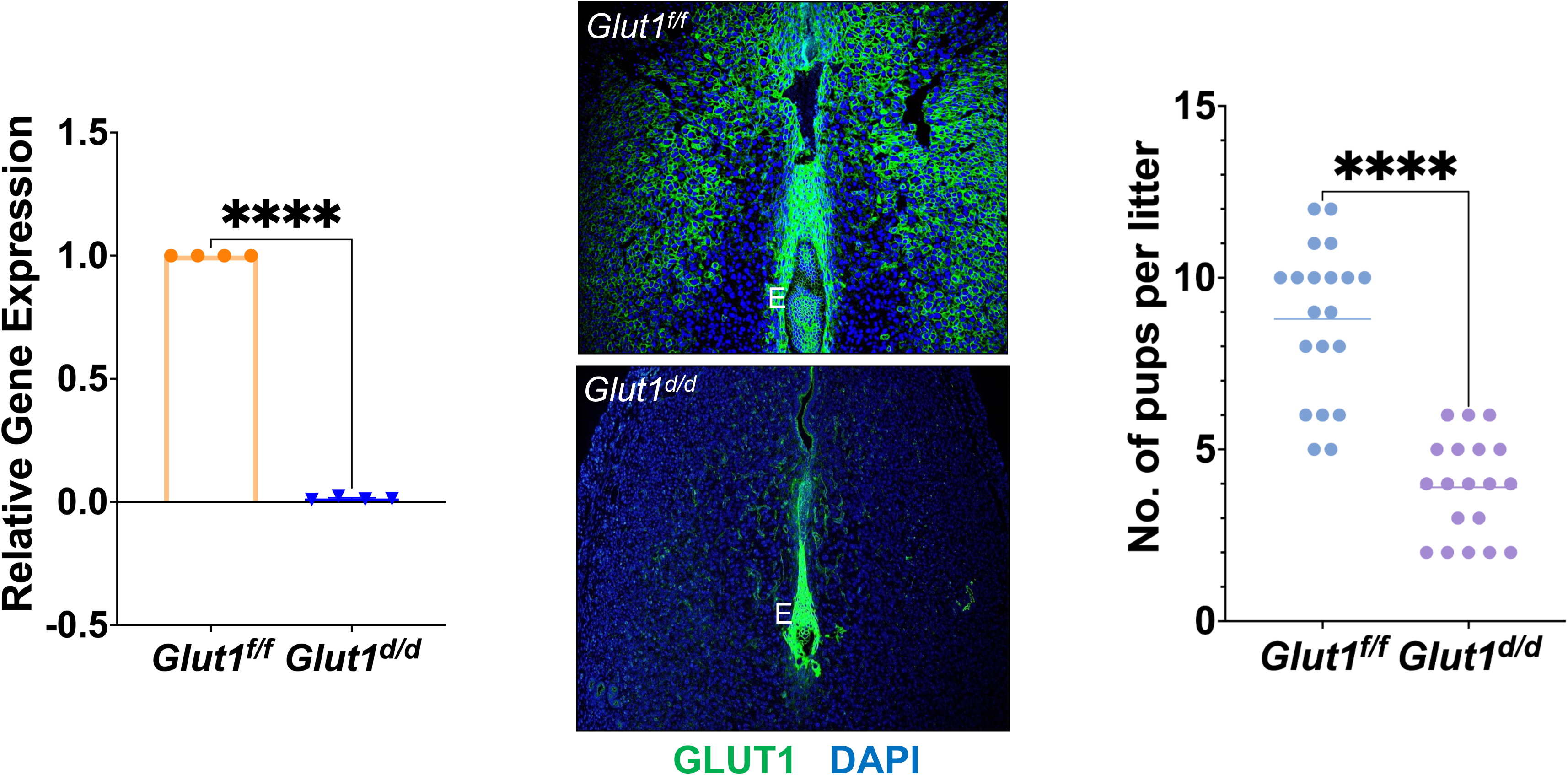
Loss of Glut1 expression in the uterus of *Glut1^d/d^* mice. **Left:** Uterine RNA was purified from *Glut1^f/f^*and *Glut1^d/d^* mice on day 7 of pregnancy and analyzed by qPCR. Relative levels of Glut1 mRNA expression in the uteri of *Glut1^d/d^* mice are compared to those in *Glut1^f/f^* control mice. Data represent mean ± SEM from three separate samples. Asterisk indicates statistically significant differences. Middle: Uterine sections obtained from day 7 pregnant *Glut1^f/f^* (top panel) and *Glut1^d/d^* (bottom panel) mice were subjected to IF using anti-Glut1 antibody. Note the lack of GLUT1 immunostaining in stromal cells of mutant mice; GLUT1 staining is retained in the implanting embryos of mutant mice. E indicates embryo. **Right:** Ablation of uterine Glut1 leads to severe female infertility. Results of a six-month breeding study are shown.

A six-month breeding study was performed by mating *Glut1^f/f^* or *Glut1^d/d^* females with wild-type males of proven fertility (Fig. 3, right). At the conclusion of the study, the total number of pups born to *Glut1^d/d^*females was reduced by more than 50% relative to *Glut1^f/f^* controls (Fig. 3, right). These data establish that the severe fertility defect of *Glut1^d/d^* females is attributable to loss of Glut1 in PGR-expressing uterine cells.

### Embryo attachment and the early stages of stromal differentiation are unaffected in Glut1^d/d^ mice

We next asked whether the subfertility of *Glut1^d/d^* females arose from a defect in embryo implantation. Gross examination of uterine morphology revealed comparable numbers of implantation sites in *Glut1^f/f^*and *Glut1^d/d^* females on day 7 of pregnancy (Fig. 4A). Histological analysis, however, showed that embryos implanting within the decidual bed of *Glut1^d/d^* uteri were noticeably smaller than those in controls (Fig. 4B), prompting a closer examination of decidualization.

**Figure 4.**
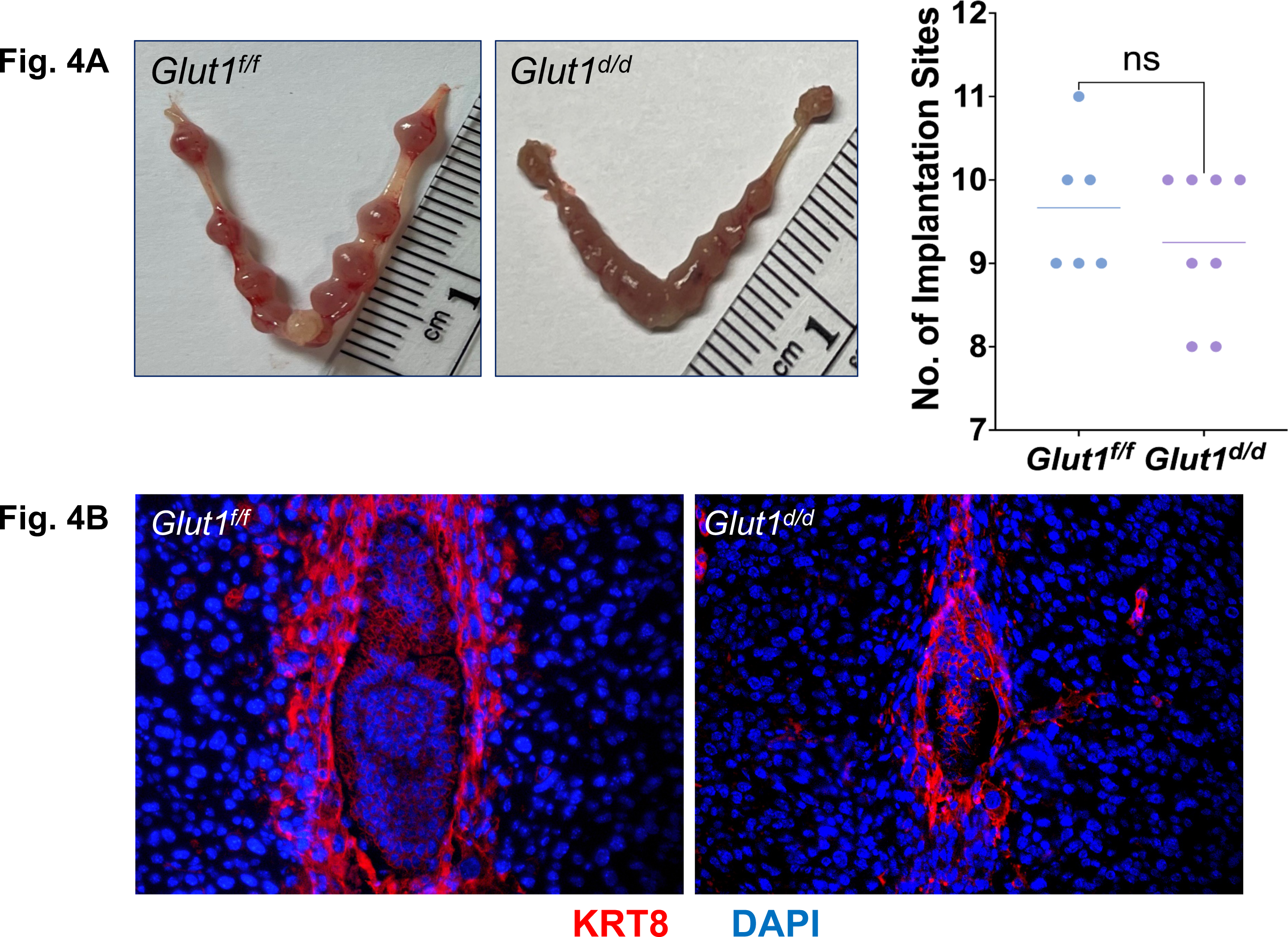
Early implantation is unaffected in the Glut1 conditional knockout mouse. **A.** Gross morphology of *Glut ^f/f^* and *Glut1^d/d^* uteri at day 7 of gestation. Note the comparable number of implantation sites in the two genotypes. **B.** Uterine sections obtained from day 7 pregnant *Glut1^f/f^* and *Glut1^d/d^* mice were nh subjected to IF using an anti-cytokeratin antibody. Blue: DAPI. Data were repeated with N=3 mice. Representative images are shown.

Decidualization involves stromal proliferation followed by differentiation into decidual cells. Stromal proliferation, assessed by phospho-histone H3 staining, was comparable between *Glut1^f/f^* and *Glut1^d/d^* uteri on day 6 of pregnancy (Fig. 5A). Likewise, alkaline phosphatase staining, a known biomarker of stromal differentiation, revealed no overt difference between genotypes on day 7 (Fig. 5B, left). Expression of the canonical decidualization regulators *Esr1*, *Pgr*, *Cebpb*, and *Hand2* was similarly unaffected in *Glut1^d/d^* uteri (Fig. 5B, right). In contrast, expression of *Prl8a2*, a late marker of differentiation, and *Cx43*, a gap junction protein, was significantly reduced in day 7 *Glut1^d/d^* uteri, indicating that selective late-stage features of decidualization are compromised by loss of stromal *Glut1*. Of note, expression of *Glut4* in stromal cells was unaffected (Fig. 5B), consistent with a non-redundant role for *Glut1*.

**Figure 5.**
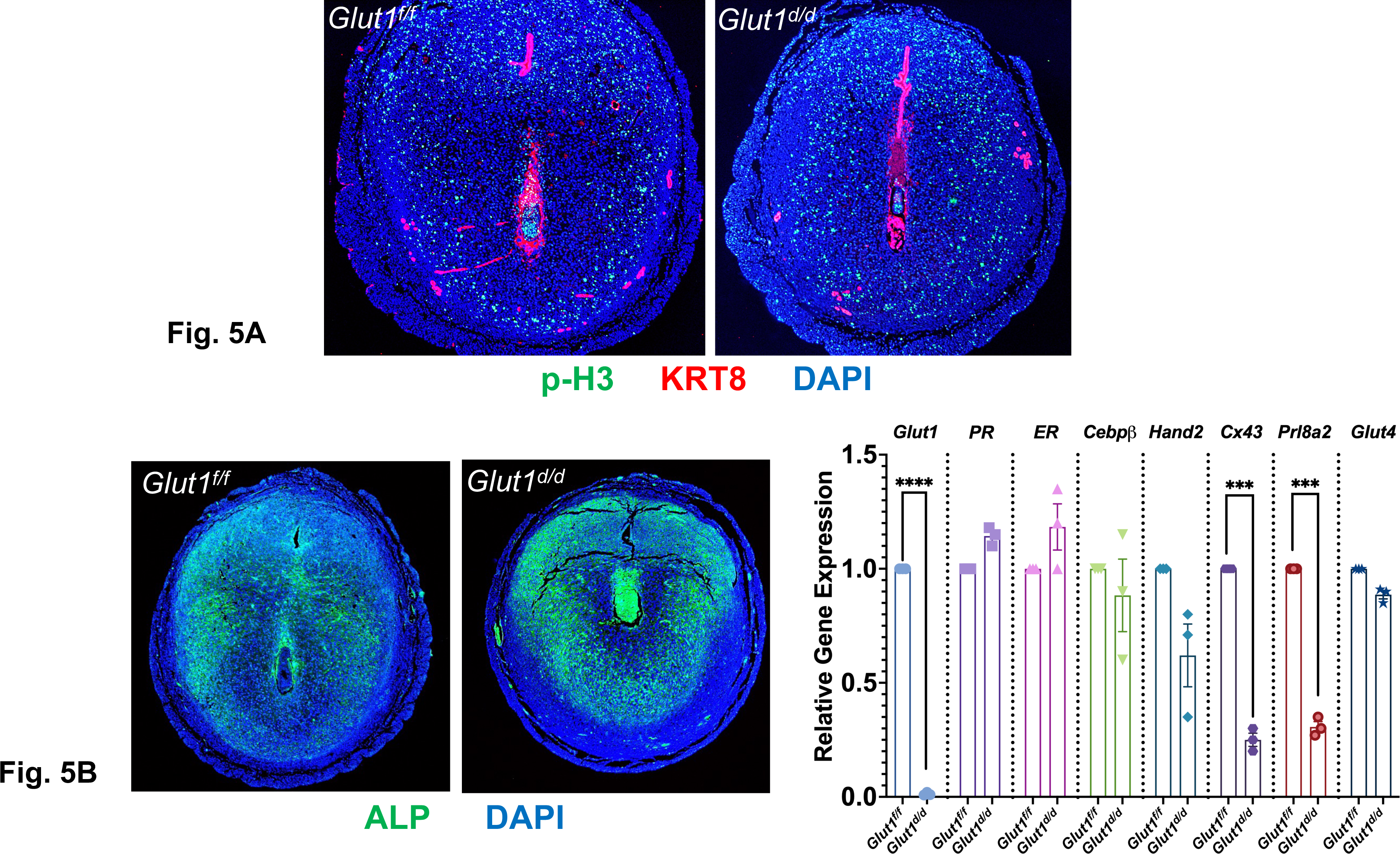
Assessment of decidualization in the Glut1 conditional knockout mouse. **A.** Uterine sections obtained from day 6 pregnant *Glut1^f/f^* and *Glut1^d/d^* mice were subjected to IF using anti-phospho histone H3 antibody to assess cell proliferation (N=3). **B. Left:** Uterine sections from *Glut1^f/f^* and *Glut1^d/d^*mice on day 7 of pregnancy were subjected to IF using an antibody against alkaline phosphatase, an early marker of stromal cell differentiation (N=3). **Right:** Total RNA was isolated from uteri on day 7 of pregnancy, and qPCR analysis was performed using primers specific for *Glut1*, *Pgr*, *Esr1*, *Cebpb*, *Hand2*, *Cx43*, *Prl8a2*, and *Glut4*.

### Glut1-Mlx maintains a Hif2α-Rab27b feed-forward circuit during decidualization

To gain insight into the pathways perturbed by disrupted glucose homeostasis in *Glut1*-null uteri, we performed RNA-sequencing on day 7 pregnant *Glut1^f/f^* and *Glut1^d/d^* uteri. Strikingly, expression of Hif2α and its downstream effector Rab27b was significantly reduced in *Glut1^d/d^* uteri (Fig. 6A, left). To corroborate this finding at the protein level, we performed immunocytochemistry for HIF2α in endometrial stromal cells. Consistent with the RNA profile, robust HIF2α protein expression in *Glut1^f/f^* stromal cells was sharply diminished in *Glut1^d/d^* cells (Fig. 6A, right), establishing that Glut1 sustains Hif2α expression during decidualization.

**Figure 6.**
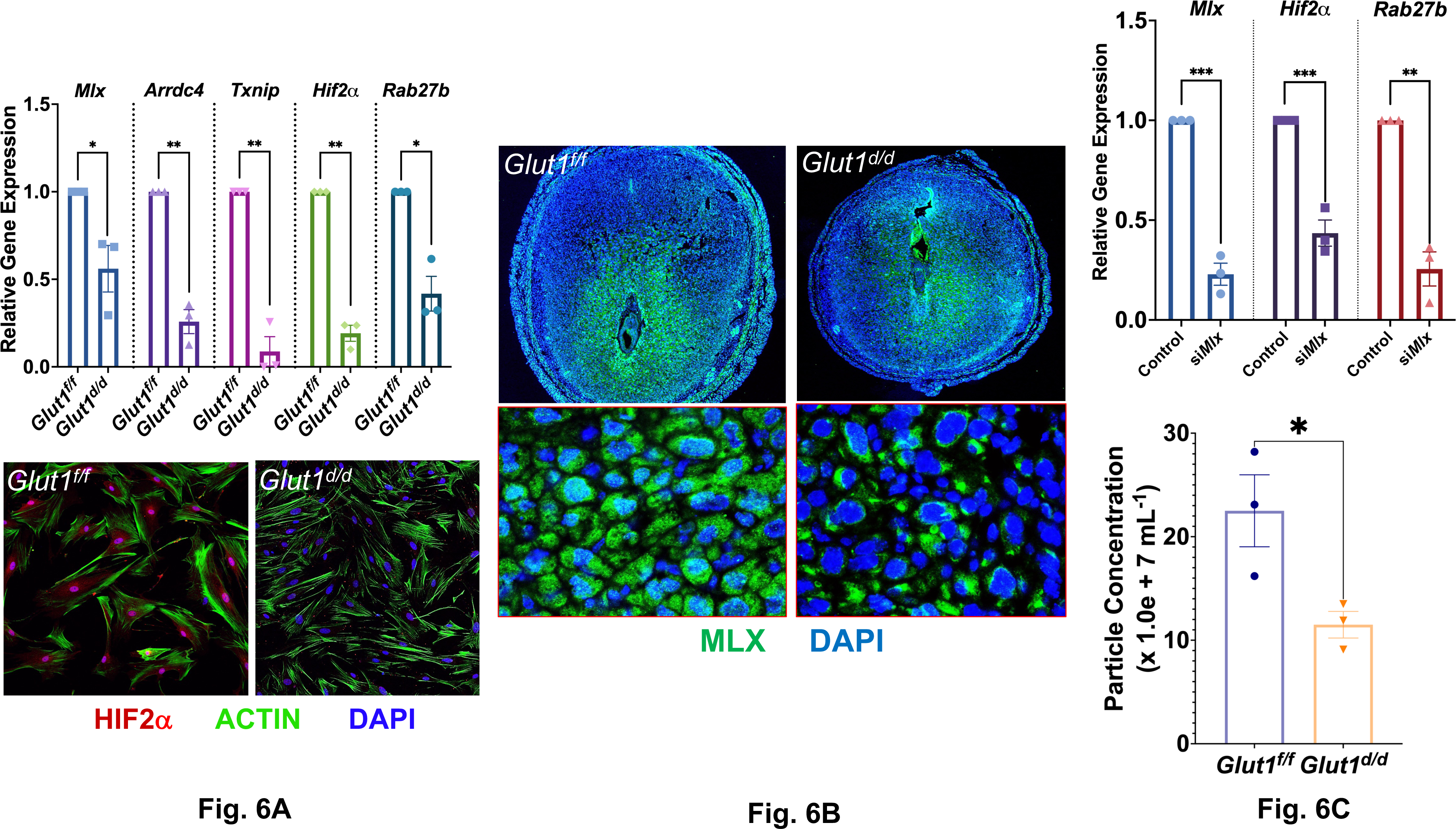
Glut1-Mlx pathway regulates Hif2α-Rab27b in the uterus during decidualization. **A. Left:** qPCR was performed to analyze the expression of *Mlx*, *Arrdc4*, *Txnip*, *Hif2α*, and *Rab27b* in the uteri of *Glut1^f/f^* and *Glut1^d/d^* mice on day 7 of pregnancy. Data represent mean ± SEM from four separate samples and were analyzed by t-test. Asterisks indicate statistically significant differences (*P < 0.05, **P < 0.01). **Right:** Mouse endometrial stromal cells isolated from *Glut1^f/f^* and *Glut1^d/d^* uteri on day 4 of pregnancy were cultured for 48 h, fixed, and subjected to IF using an HIF2α antibody. Data were repeated with N=3 mice; representative images are shown. **B.** Uterine sections obtained from day 7 pregnant Glut1f/f and *Glut1^d/d^*mice were subjected to IF using an anti-MLX antibody. Lower images show magnified views of the boxed areas in the upper panels. **C. Left:** MESCs were transfected with Mlx-specific siRNA or scrambled siRNA controls. Cells were lysed, total RNA was isolated, and real-time PCR was performed for *Mlx*, *Hif2α*, and *Rab27b*. Relative gene expression was calculated by setting the scrambled siRNA-treated sample to 1.0. RPLP0 (36B4) was used to normalize RNA levels. Data were collected from three independent samples. **Right:** MESCs isolated from the uteri of day 4 pregnant mice were subjected to in vitro decidualization. EVs were isolated from conditioned media at 48 h and analyzed by MRPS. Data are mean fold changes ± SEM.

We also observed a marked decline in transcripts encoding the Max-like protein X (MLX), a glucose-sensing transcription factor (15–17), and its downstream targets, Arrdc4 and Txnip, in *Glut1^d/d^*uteri (Fig. 6A, left). MLX protein was correspondingly reduced (Fig. 6B), and importantly, the residual MLX in *Glut1^d/d^* uteri was excluded from the nucleus, in contrast to the predominantly nuclear MLX of *Glut1^f/f^* uteri (Fig. 6B). This redistribution is consistent with the established mechanism in which MLX, upon sensing intracellular glucose, translocates from cytoplasm to nucleus to bind carbohydrate response elements and regulate transcription (15–17). This lack of nuclear MLX protein in decidual cells indicated an impairment of glucose metabolic signaling in *Glut1^d/d^* uteri.

These observations led us to consider that, while Hif2α regulates Glut1 at the time of implantation, Glut1 may, in turn, regulate Hif2α during decidualization, a feed-forward loop that would sustain Hif2α expression as the decidualization progresses. To test whether Mlx, acting downstream of Glut1, is required for this control, we performed siRNA-mediated knockdown of *Mlx* in mouse endometrial stromal cells (MESCs) isolated from day 4 pregnant uteri (Fig. 6C, left). Knockdown of *Mlx* produced a marked downregulation of *Hif2α* and *Rab27b*. Because we previously showed that the Hif2α-Rab27b axis controls extracellular vesicle (EV) secretion as a mediator of cell-cell communication during decidualization (18–20), the loss of this axis in *Glut1^d/d^* cells suggested that EV secretion would be impaired.

To test this, MESCs from *Glut1^f/f^* and *Glut1^d/d^*day 4 uteri were subjected to in vitro decidualization, and EVs were harvested from conditioned media at 72 h and quantified by microfluidic resistive pulse sensing (MRPS). EV secretion by *Glut1^d/d^* stromal cells was significantly reduced relative to controls (Fig. 6C, right). Together, these data identify a Glut1-Mlx-Hif2α-Rab27b axis through which stromal glucose sensing sustains EV-mediated cell-to-cell communication in the uterus during decidualization.

### Decidual angiogenesis is impaired in Glut1^d/d^ mice

To determine whether Glut1 deficiency alters the molecular cargo of EVs, we performed comparative mass spectrometry on EVs isolated from in vitro stromal cultures derived from *Glut1^f/f^* and *Glut1^d/d^*mice. Quantitative proteomic analysis, as described in Materials and Methods, identified a set of differentially abundant proteins in *Glut1^d/d^* EVs that segregate into two functionally related groups: regulators of decidual angiogenesis and regulators of trophoblast differentiation (Tables 1A and 1B).

Among angiogenesis regulators, *Glut1^d/d^* EVs were enriched in molecules that ordinarily restrain endothelial sprouting and capillary morphogenesis (Table 1A). Thrombospondin-1 (Thbs1), an inhibitor of endothelial proliferation, migration, and tube formation (43), was elevated more than five-fold; Reelin (Reln), a secreted glycoprotein with reported activity on endothelial behavior, was elevated nearly 13-fold, and CCN2/CTGF, a context-dependent matricellular regulator of vascular morphogenesis (44) together with the basement-membrane-binding glycoprotein extracellular matrix protein 1 (Ecm1) were each markedly increased. In striking contrast, two factors that promote vascular and matrix remodeling, type XV collagen (Col15a1) and matrix metallopeptidase 19 (Mmp19), an ECM remodeler (45), were sharply depleted from *Glut1^d/d^* EVs (>10-fold reduction). This composite signature of upregulated antiangiogenic and downregulated pro-remodeling cargoes predicts a paracrine environment unfavorable to decidual vascular development.

**Table 1.**
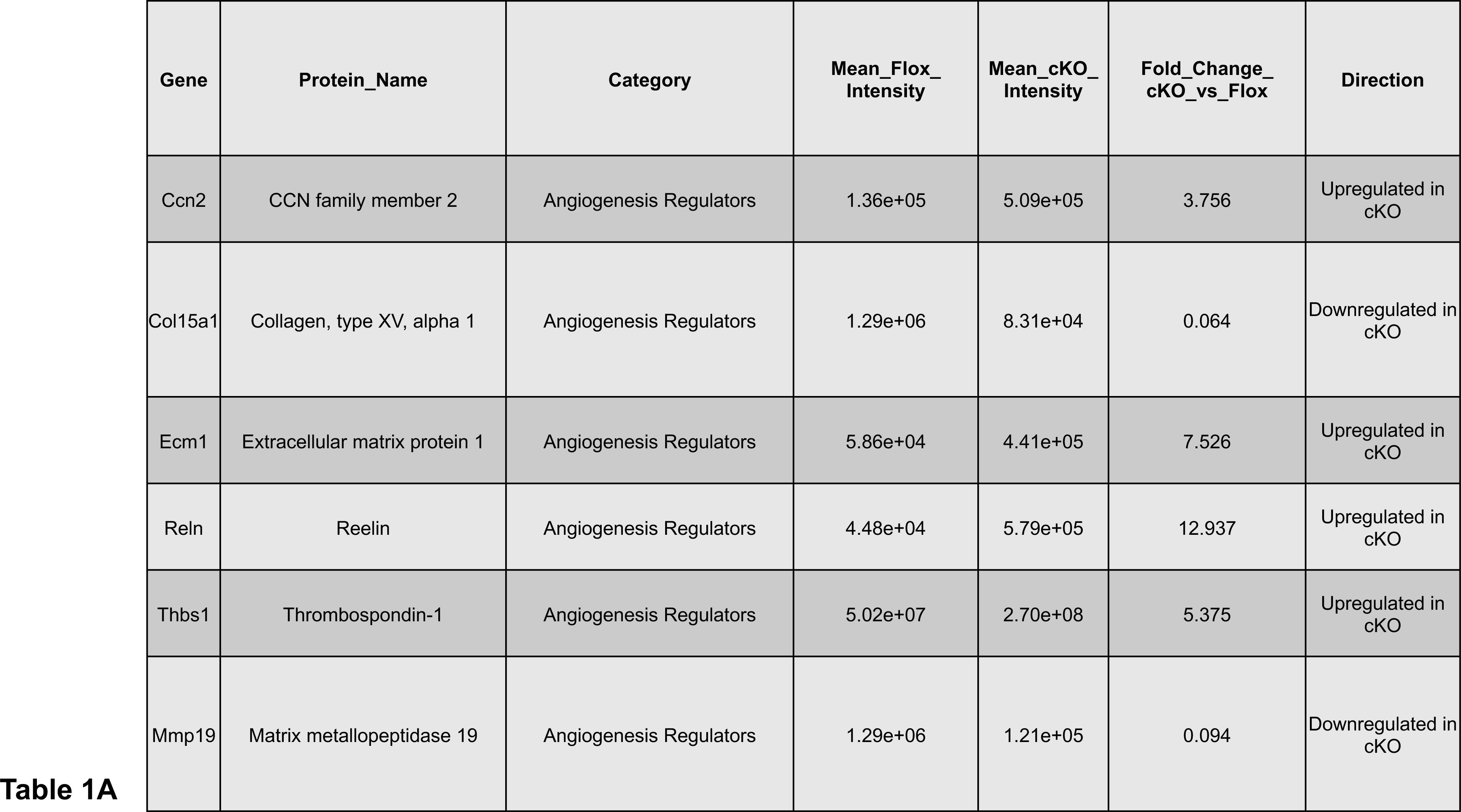
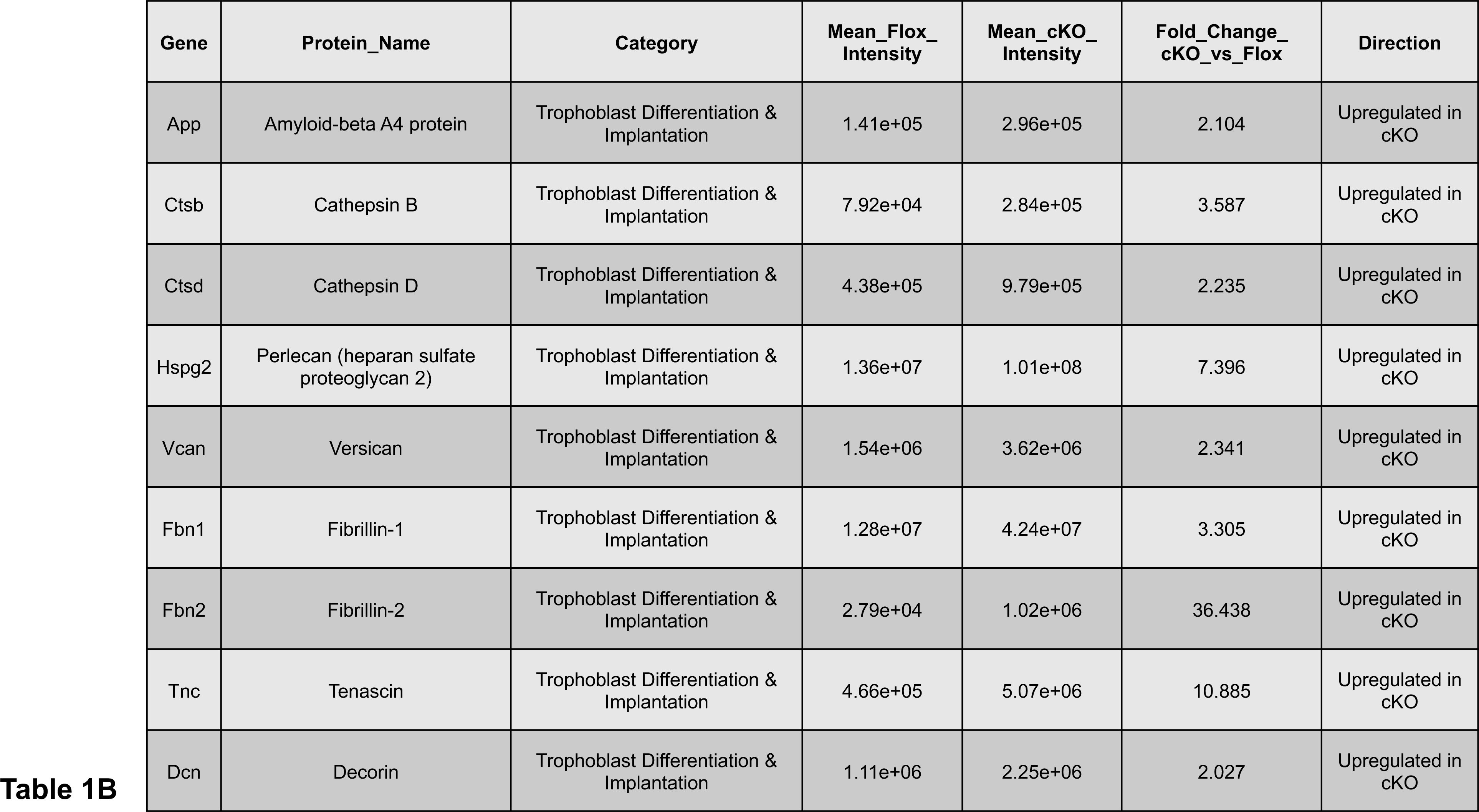
A partial list of EV cargoes involved in angiogenesis (A) and trophoblast differentiation (B) secreted during decidualization of MESCs collected from uteri of *Glut1^f/f^*(Flox) and *Glut1^d/d^* (cKO) mice on day 7 of pregnancy, followed by 72 h of culture in vitro.

To determine whether these alterations in EV cargo translated into functional defects *in vivo*, we next examined the establishment of the decidual vascular network during early pregnancy. In *Glut1^f/f^*mice on day 7 of pregnancy, immunostaining for platelet/endothelial cell adhesion molecule 1 (PECAM1) revealed a well-developed vascular network throughout the decidual bed surrounding the implanted embryo (Fig. 7, left). In contrast, PECAM1 staining was markedly reduced in *Glut1^d/d^*uteri (Fig. 7, right). Notably, PECAM1-positive cells were present in *Glut1^d/d^* uterine sections but failed to sprout from the stump (white square, Fig. 7A, right) and form an angiogenic network, consistent with impaired vascular development in the absence of stromal Glut1.

**Figure 7.**
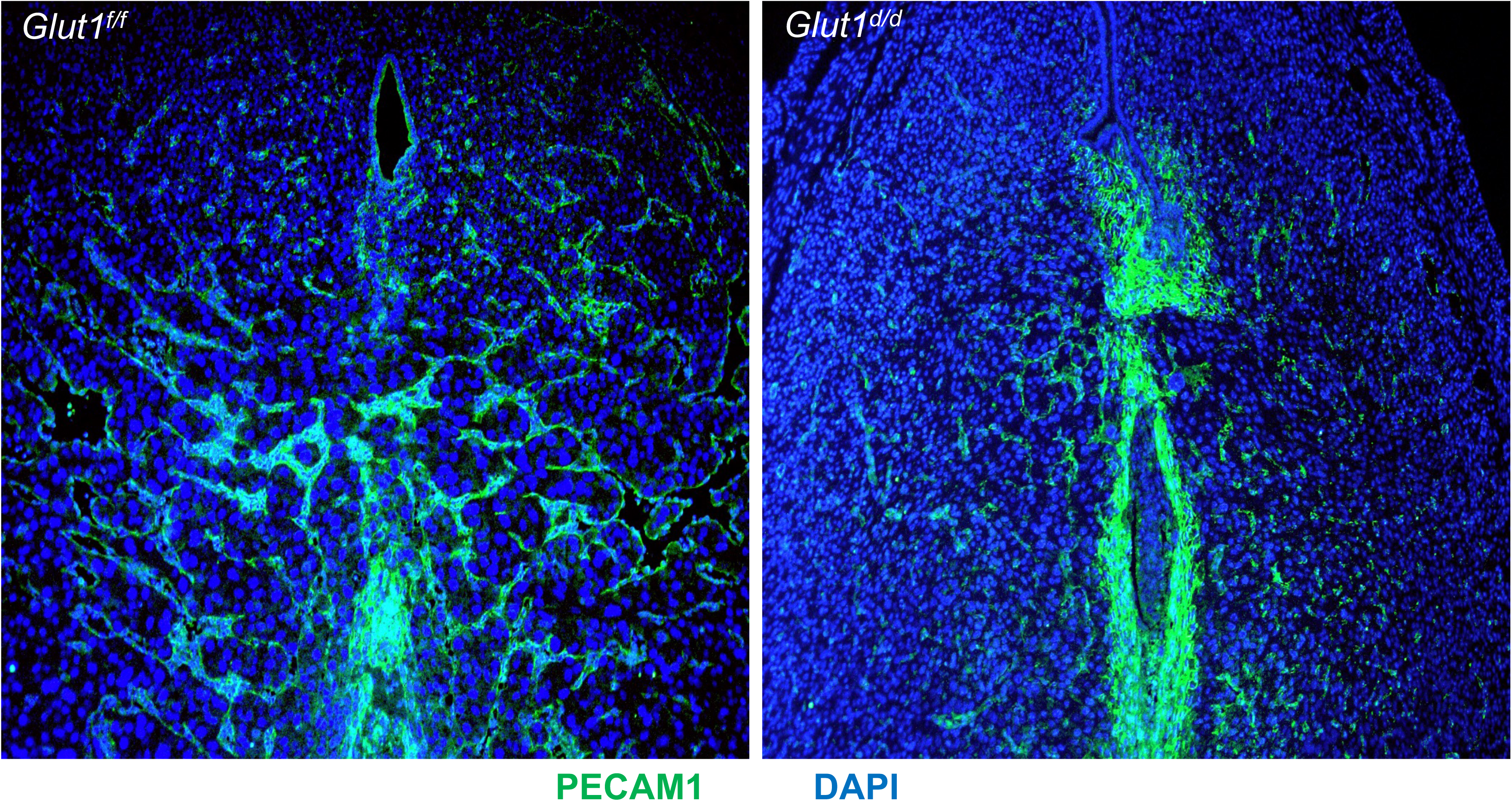
Endometrial stromal angiogenesis is impaired following Glut1 loss in the uterus. Uterine sections from *Glut1^f/f^* and *Glut1^d/d^* mice on day 7 of pregnancy were subjected to IF using a PECAM1 (red) antibody. N=3 mice were examined for each genotype.

### Trophoblast differentiation and placentation are dysregulated in Glut1^d/d^ uteri

A second cluster of dysregulated EV cargoes comprised regulators of trophoblast invasion, differentiation, and ECM signaling (Table 1B). Cathepsin B and cathepsin D (Ctsb, Ctsd), lysosomal proteases required for trophoblast invasion and trophoblast giant cell differentiation (45, 46), were both elevated in *Glut1^d/d^* EVs. The basement-membrane heparan sulfate proteoglycan perlecan (Hspg2) and the small leucine-rich proteoglycan decorin (Dcn), a known suppressor of trophoblast migration in disorders such as preeclampsia (47), were similarly upregulated. Additional cargoes included the ECM-associated proteins versican (Vcan), fibrillin-1 (Fbn1), fibrillin-2 (Fbn2), tenascin-C (Tnc), and amyloid-β A4 protein (App), factors that collectively contribute to extracellular matrix remodeling and regulation of trophoblast behavior at the maternal-fetal interface (46). Collectively, this altered EV cargo profile predicts a decidual microenvironment that is unsupportive for trophoblast differentiation and placental development.

To determine whether these molecular alterations translate into defects in trophoblast lineage progression in vivo, we next examined trophoblast differentiation at gestational day 8. IF analysis for PCNA, a proliferation marker, and KRT8, a trophoblast marker, within the ectoplacental cone (EPC) revealed a striking expansion of PCNA-positive trophoblast cells in *Glut1^d/d^*females compared with controls (Fig. 8). Under normal conditions, trophoblast progenitors undergo a tightly regulated transition from proliferation to differentiation during placental development. The persistence and expansion of proliferative trophoblast cells in *Glut1^d/d^* implantation sites suggest that loss of Glut1 disrupts this developmental transition, leading to sustained proliferation of trophoblast progenitors and compromising differentiation.

**Figure 8.**
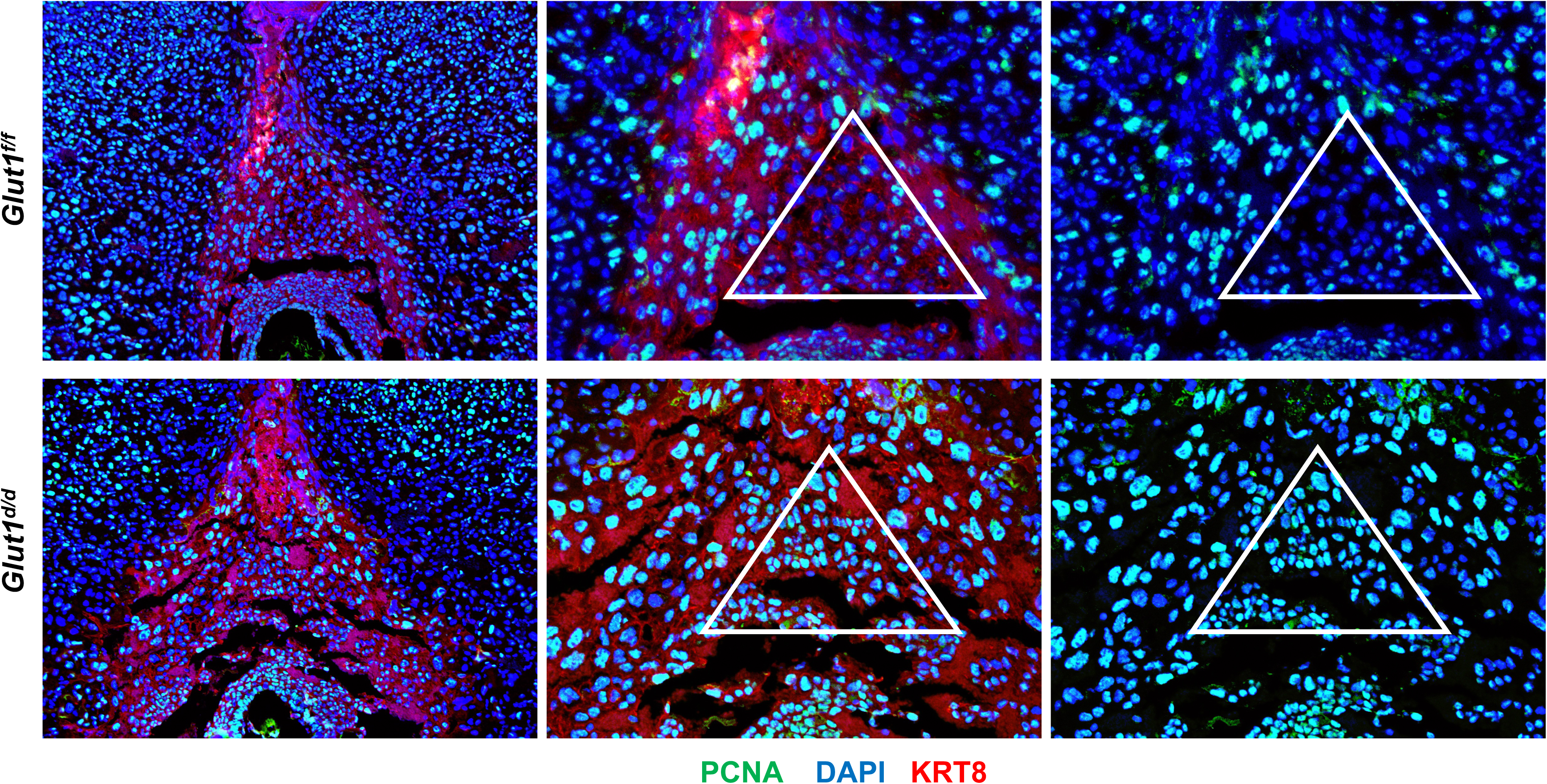
Trophoblast differentiation is altered following endometrial Glut1 loss. Uterine sections from *Glut1^f/f^* and *Glut1^d/d^* mice on day 8 of pregnancy were subjected to IF using PCNA, KRT8, and DAPI. The encircled area shows the trophoblast progenitor cells in the ectoplacental cone (EPC). N=3 mice were examined for each genotype.

Consistent with these early defects in trophoblast differentiation, placental development progressively deteriorated as gestation advanced. Although no overt loss of implanted embryos was apparent in *Glut1^d/d^*uteri through day 7 of gestation, embryos undergoing resorption became evident starting on day 10 (Fig. 9, left). By day 12, fetal-placental units in *Glut1^d/d^* uteri exhibited markedly smaller placentas and fetuses than *Glut1^f/f^* controls (Fig. 9, right). To further assess placental development, we performed IF on day 10 placental sections using an antibody against placental lactogen 1 (PL1) (Fig. 10A). PL1 is normally expressed in trophoblast giant cells (TGCs) on day 9, peaks on day 10, and declines by day 12 (21–23). *Glut1^f/f^* placentas displayed PL1 expression in 1-2 layers of trophoblast cells on day 10 that declined by day 12 (Fig. 10A, upper). In striking contrast, *Glut1^d/d^* placentas exhibited multiple layers of TGCs on days 10-12 of pregnancy (Fig. 10A, lower), consistent with aberrant expansion of this trophoblast population.

**Figure 9.**
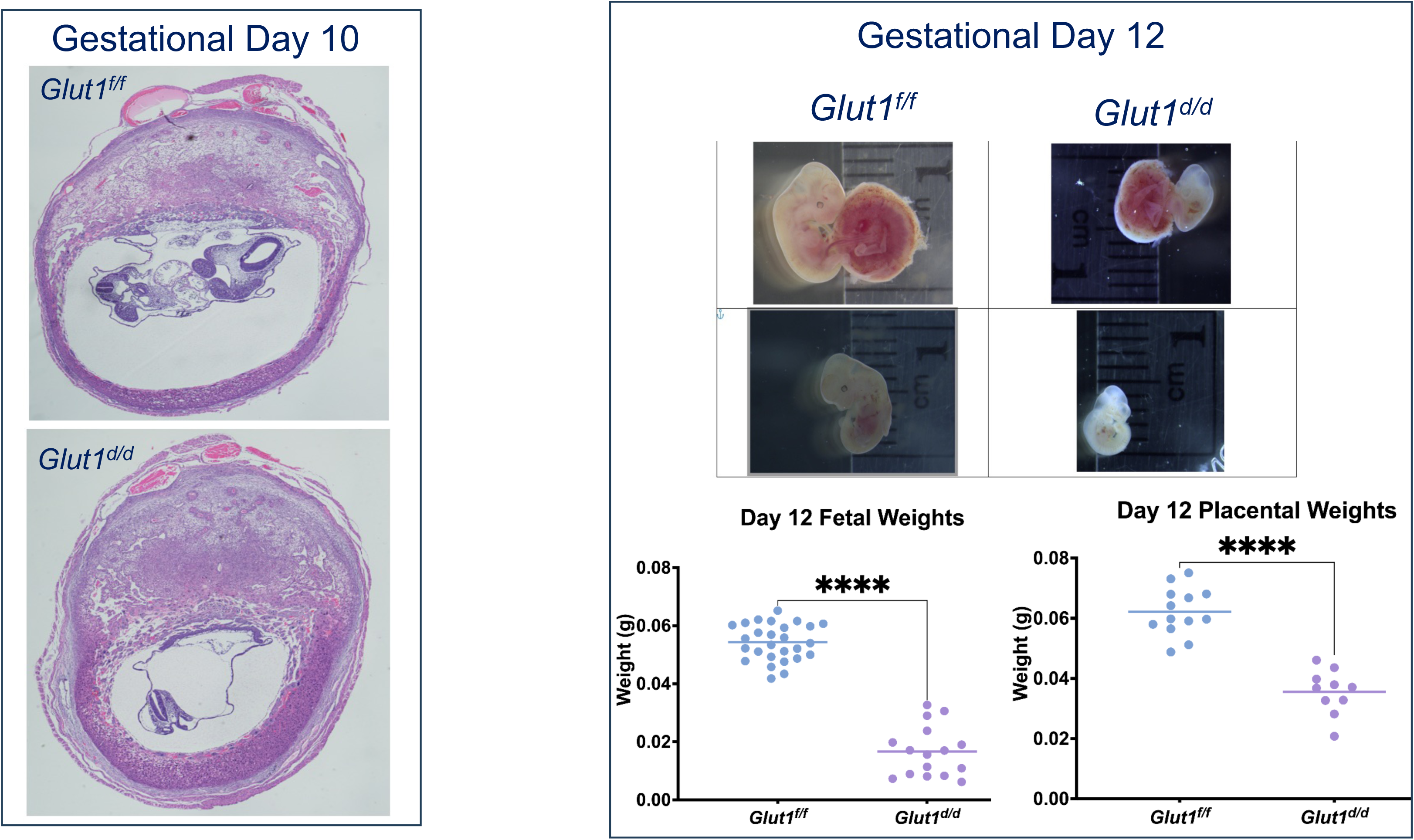
Pregnancy failure in Glut1-null mice in mid-gestation. **Left:** H&E staining of uteroplacental sections from *Glut1^f/f^* and *Glut1^d/d^* mice on day 10 of pregnancy. **Right:** Gross morphology of *Glut1^f/f^* and *Glut1^d/d^* mice on day 12 of gestation. Dissection of implantation sites revealed underdeveloped placentas and embryos in *Glut1^d/d^* mice. Fetal and placental weights are shown in the lower panels. N=4/genotype on each day.

**Figure 10.**
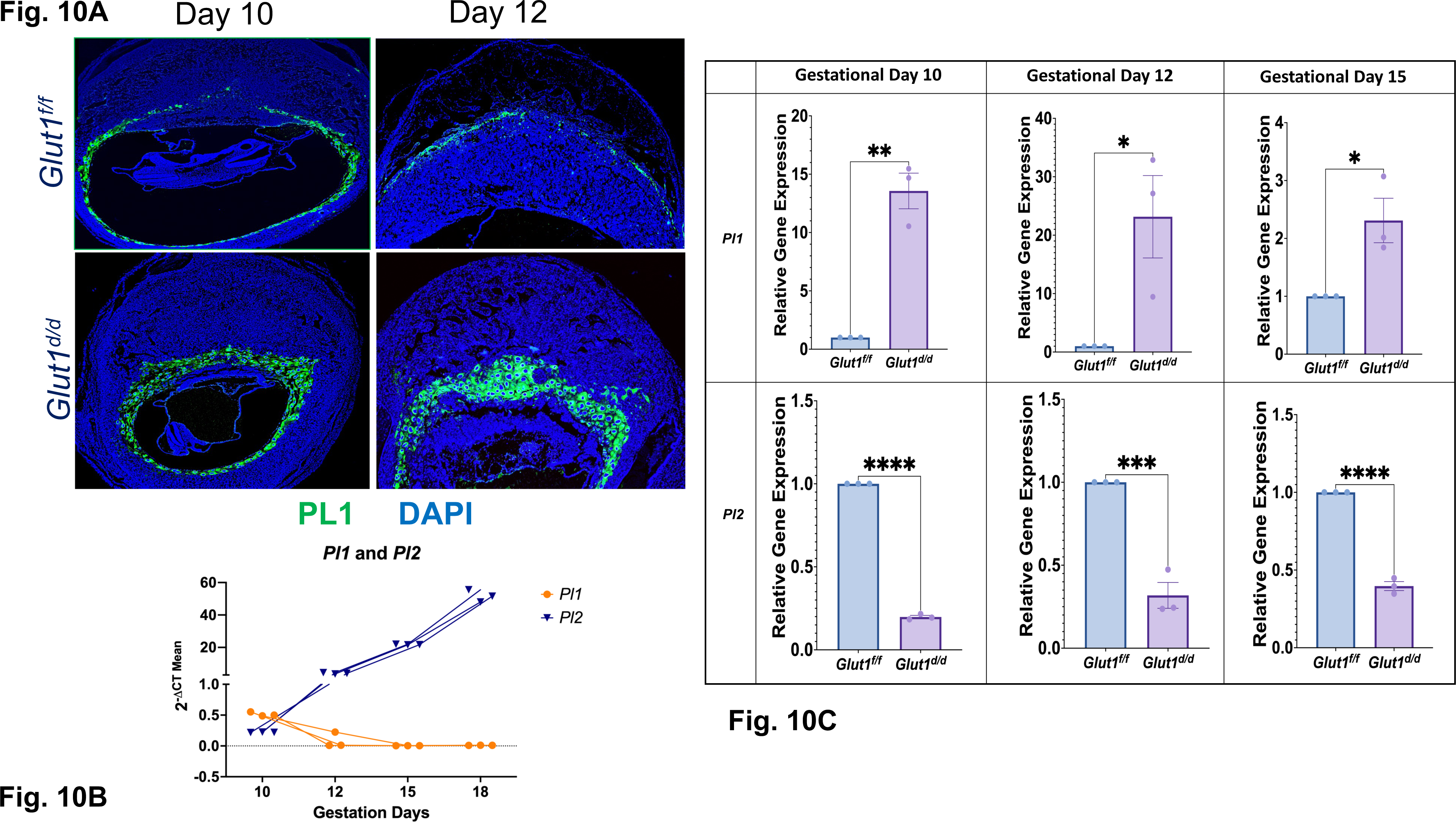
Abnormal trophoblast differentiation in *Glut1^d/d^*. **A.** Uterine sections from *Glut1^f/f^* and *Glut1^d/d^* mice on days 10 and 12 of pregnancy were subjected to IF using an antibody against PL1. Data are representative images from N=3. **B.** Placental gene expression (2^-ΔCT^ ^Mean^) of *Pl1* (orange circles) and *Pl2* (blue triangles) quantified in control (*Glut1^f/f^*) mice across gestational days 10, 12, 15, and 18. Individual biological replicates are plotted with connecting trendlines to illustrate the progressive downregulation of *Pl1* concurrent with the robust upregulation of *Pl2* over time. **C.** Total RNA was isolated from the placenta on D10, D12, and D15 of pregnancy, and qPCR analysis was performed using primers specific for *Pl1* and *Pl2*. Data represent mean ± SEM from three separate samples. Asterisks indicate statistically significant differences (*P<0.05, **P<0.01, ***P<0.001, ****P<0.0001).

To determine whether this altered trophoblast differentiation program extended beyond PL1-positive TGCs, we next examined the expression of placental lactogen 2 (PL2), a marker of later trophoblast differentiation associated with spongiotrophoblasts in the junctional zone. Whereas PL1 expression normally declines by day 12, PL2 levels typically increase beginning on day 11 and remain elevated throughout gestation (21, 24). Indeed, during normal gestation in control (*Glut1^f/f^*) mice, placental expression of *Pl1* and *Pl2* exhibited distinct, opposing temporal profiles, characterized by a progressive decline in *Pl1* levels alongside a robust, increase in *Pl2* expression from gestational day 10 through day 18 (Fig. 10B). Interestingly, qPCR analysis of placental lactogens in *Glut1^f/f^* and *Glut1^d/d^* placentas revealed a striking dysregulation of expression kinetics of *Pl1* and *Pl2* throughout gestation (Fig. 10C). While *Pl1* transcripts remained elevated in *Glut1^d/d^* uteri relative to controls through gestational day 15, *Pl2* transcripts stayed significantly below control levels over the same window (Fig. 10C). Together, these data indicate that endometrial Glut1 loss causes a sustained shift in the temporal program of trophoblast-derived endocrine factors, with failure of the PL1-to-PL2 transition that normally accompanies trophoblast differentiation.

Given the expansion of PCNA-positive progenitors within the EPC (Fig. 8), we next asked whether this aberrant proliferation altered late-gestational placental architecture. Specifically, we investigated the lineage commitment of trophoblast cells within the junctional zone (JZ) at day 15. Periodic Acid–Schiff–Hematoxylin (PAS-H) staining was used to quantify glycogen trophoblast (GlyT) cells, which originate from EPC progenitors. As shown in Fig. 11, *Glut1^d/d^* placentas exhibited a striking accumulation of GlyT cells relative to *Glut1^f/f^* controls, suggesting that Glut1 is required to maintain the balance of trophoblast subtypes. Notably, a similar expansion of the junctional zone (JZ) and increased GlyT cell abundance have been reported in metabolic disorders such as Gestational Diabetes Mellitus (GDM) (34, 41, 42).

**Figure 11.**
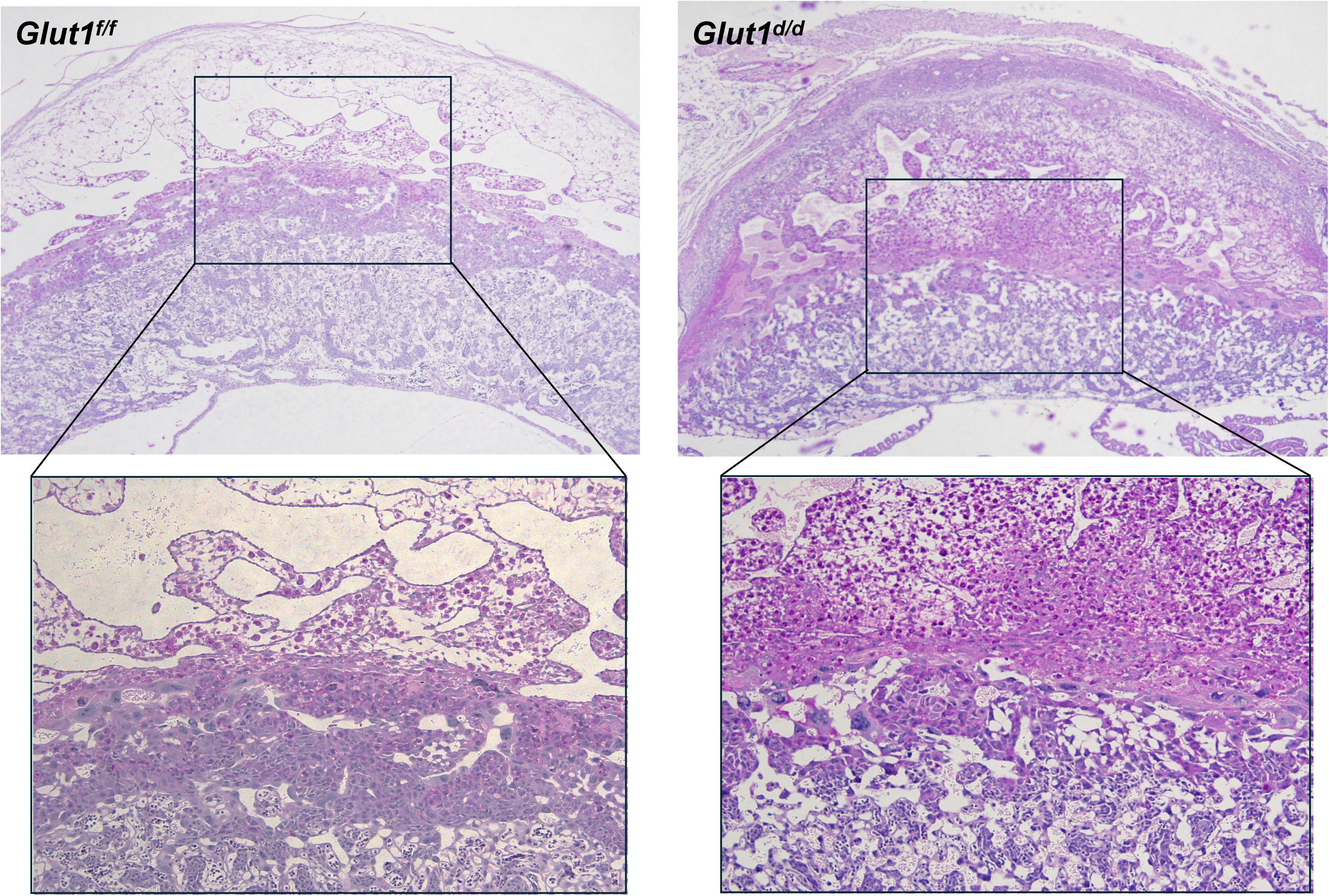
Increase in the glycogen trophoblast (GlyT) population in the Glut1^d/d^ mice. PAS-H staining of uteroplacental sections from *Glut1^f/f^* and *Glut1^d/d^* mice on day 15 of pregnancy.

### Glut1^d/d^ mice spontaneously develop a gestational diabetes phenotype

Interestingly, PL2 is a major lactogenic hormone in the maternal serum throughout gestation (21) and plays an important role in maternal metabolism and glucose homeostasis. It can bind to the prolactin receptor on pancreatic β-cells, promoting β-cell proliferation and insulin secretion (25). Pregnancy is accompanied by dynamic metabolic adaptations that ensure adequate nutrient availability for the growing fetus while maintaining maternal glucose homeostasis. To meet these increased metabolic demands, maternal pancreatic β-cells undergo functional adaptations that enhance insulin secretion during pregnancy. Failure of this adaptive β-cell response can disrupt glucose homeostasis and predispose to gestational diabetes. Because *Pl2* levels were significantly reduced in Glut1 mutants, we asked whether these mice display features of GDM. We measured blood glucose levels in *Glut1^f/f^* and *Glut1^d/d^* females before, during, and after pregnancy (Fig. 12, left). Blood glucose levels in nonpregnant mice and during early pregnancy (day 7) were comparable between the two genotypes. Strikingly, beginning on day 12, blood glucose levels were significantly elevated in *Glut1^d/d^* females relative to controls, and this trend persisted on days 15 and 18 of pregnancy. Following delivery, by day 2 postpartum, blood glucose in *Glut1^d/d^* mice had returned to *Glut1^f/f^* levels (Fig. 12, left). Consistent with these observations, insulin levels were significantly reduced in *Glut1^d/d^*females relative to controls on day 18 of pregnancy (Fig. 12, right). Together, these data demonstrate that mice lacking uterine Glut1 exhibit a sustained elevation in systemic blood glucose from mid-gestation through term, the defining clinical feature of gestational diabetes mellitus.

**Figure 12.**
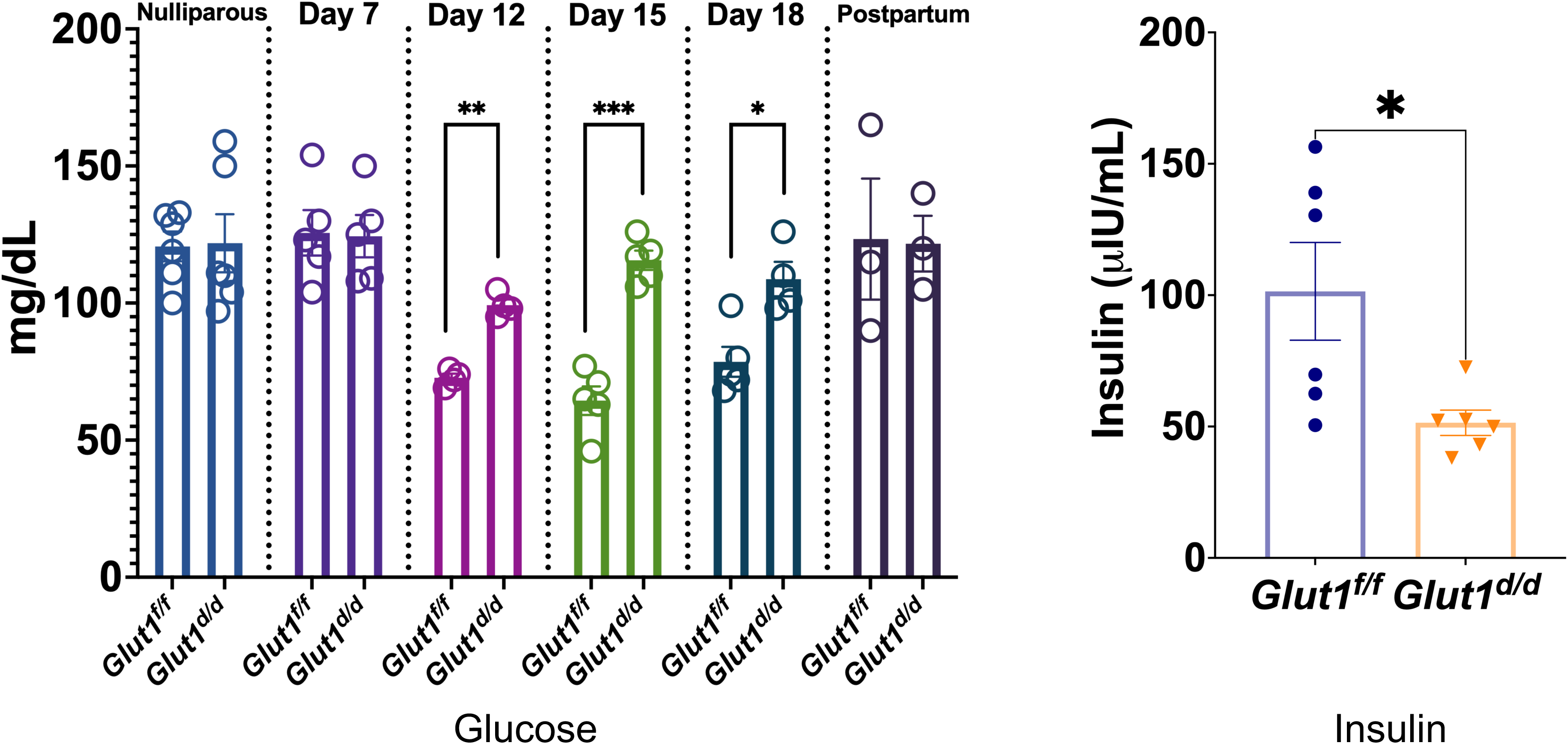
*Glut1^d/d^* mice spontaneously develop a gestational diabetes phenotype. **Left:** *Glut1^f/f^* and *Glut1^d/d^* mice at the nonpregnant stage, pregnancy days 7, 12, 15, 18, and 2 days after delivery were subjected to blood glucose measurements. **Right:** Blood insulin levels in *Glut1^f/f^* and *Glut1^d/d^* mice are shown on day 18 of pregnancy. Data represent mean ± SEM from four separate samples. Asterisks indicate statistically significant differences (*P<0.05, **P<0.01, ***P<0.001).

## DISCUSSION

Embryo implantation and the establishment of pregnancy constitute a uniquely demanding metabolic feat, with glucose serving as the dominant energy source for the periimplantation uterus (26). The differentiation of fibroblastic endometrial stromal cells into decidual cells is a critical, glucose-dependent transformation (2–4, 39); both insufficient and excessive glucose metabolism impair decidualization in human and mouse models (12), implying that endometrial glucose uptake must be carefully tuned for successful pregnancy. Yet how decidual cells acquire and channel glucose to meet the rapidly changing demands of pregnancy has remained poorly defined. Here, we identify Glut1 expressed in endometrial stromal cells as a central metabolic regulator linking uterine glucose uptake to decidual angiogenesis, trophoblast differentiation, and placentation. Strikingly, ablation of endometrial Glut1 not only causes severe subfertility but also drives a spontaneous gestational diabetes phenotype. To our knowledge, this is one of the most physiologically relevant mouse models of GDM described to date (35, 41, 42).

We recently reported that Hif2α induced in endometrial stromal cells is critical for embryo implantation and the establishment of pregnancy (9). Glut1 is induced downstream of Hif2α signaling in stromal cells, suggesting that Glut1-mediated glucose influx is required for stromal proliferation and/or differentiation during pregnancy. Our results, however, reveal a more nuanced picture: stromal proliferation and the early stages of differentiation proceed normally in the absence of Glut1, while late-stage differentiation is selectively impaired. Glucose metabolism in stromal cells thus appears to be particularly critical for the late phase of decidualization. While previous studies implicated Glut1 in decidualization, those analyses were limited to *in vitro* cell culture systems and did not address the mechanism by which Glut1 supports endometrial differentiation *in vivo* (27, 28).

Depending on cell type, the late stage of differentiation alters protein trafficking, rendering cells more secretory (29). Indeed, our previous study showed that Rac1 in endometrial stromal cells controls the late stage of differentiation by enhancing the secretion of factors such as IGFBPs and VEGF (30). Concurrently, the differentiating stromal cells progressively engage in extracellular vesicle (EV) trafficking, and we recently demonstrated that the Hif2α-Rab27b axis governs EV secretion in endometrial stromal cells (18–20). Our findings place Glut1-mediated glucose uptake upstream of this axis. Glucose entry through Glut1 drives Mlx nuclear translocation and the resulting transcriptional program, which in turn sustains Hif2α and Rab27b expression. This feed-forward circuit ensures that EV-mediated cell-cell communication is sustained during decidualization. Mass spectrometry-based profiling of stromal EVs has revealed that these vesicles harbor a diverse cargo of angiogenic and trophoblast differentiation-related factors (18). Comparative proteomic analysis of EVs from *Glut1^d/d^*and *Glut1^f/f^* stromal cells identified a set of differentially abundant proteins, a large fraction of which are established regulators of decidual angiogenesis and trophoblast differentiation, providing a plausible mechanism by which Glut1 deficiency disrupts the early gestational environment.

The cargo signature itself offers two particularly informative insights into how endometrial glucose metabolism shapes the maternal-fetal interface. First, the reciprocal redistribution of pro- and antiangiogenic cargoes, such as enrichment of thrombospondin-1 (43), reelin, CCN2/CTGF (44), and ECM1, together with the depletion of basement-membrane-stabilizing type XV collagen and the matrix metalloproteinase MMP19 (45), constitutes an EV-encoded shift toward a vascularly restrictive paracrine environment and provides a plausible explanation for the failure of endothelial sprouting in *Glut1^d/d^* decidua. Second, the parallel upregulation of decorin (47), perlecan, fibrillins, versican, tenascin-C, and the lysosomal proteases cathepsin B and cathepsin D (45, 46) signals a marked rewiring of the extracellular matrix-derived cues that instruct trophoblast lineage commitment (46), and likely underlies the aberrant expansion of trophoblast progenitors, the accumulation of glycogen trophoblast cells, and the skewed PL1/PL2 endocrine output of *Glut1^d/d^* placentas. More broadly, these findings raise the possibility that the maternal metabolic state in the endometrium tunes the molecular content of stromal-derived EVs. Disruption of this metabolism-EV axis may therefore contribute to the spectrum of placenta-based pregnancy disorders, including intrauterine growth restriction, preeclampsia, and gestational diabetes (31, 41, 42).

Proper proliferation and differentiation of trophoblast cells, essential for the formation of a functional placenta, are in turn shaped by maternal factors secreted by decidual cells (40). A central phenotypic consequence of endometrial Glut1 loss is a striking abnormality in placental development. Combined with impaired decidual angiogenesis, this defect drives fetal loss and intrauterine growth restriction in *Glut1^d/d^*mice. In mammals, intrauterine growth is determined principally by nutrient delivery to the fetus via the placenta, which depends on placental size, morphology, transporter availability, and blood flow; abnormal placental development thus directly compromises fetal growth.

Detailed analysis of Glut1-null placentas revealed a marked disruption of placental architecture. Histologically, we observed an aberrant expansion of the PCNA-positive trophoblast stem cell pool within the *Glut1^d/d^* ectoplacental cone, accompanied on gestation day 15 by a marked accumulation of GlyT cells. Together, these data suggest that loss of Glut1 causes a persistent proliferative state in early progenitors that ultimately manifests as an expanded junctional-zone lineage later in gestation. Furthermore, while control mice exhibited the expected one- to two-layer TGCs expressing PL1 on day 10, multiple layers of TGCs were evident at the maternal-fetal interface in *Glut1^d/d^* mice. By day 12, the typical decline of Pl1 and the rise of Pl2 were inverted in *Glut1^d/d^* uteri. While Pl1 remained elevated, Pl2 was significantly reduced, indicating that the temporal program of trophoblast differentiation has been derailed. Aberrant trophoblast differentiation and function are at the root of many placenta-based pregnancy disorders, including intrauterine growth restriction, gestational diabetes, and preeclampsia (31, 40).

Importantly, Pl2 itself plays a key role in regulating maternal metabolism and glucose homeostasis during pregnancy. Pregnancy is physiologically associated with a gradual rise in insulin resistance, an adaptive mechanism that ensures adequate glucose supply to the rapidly growing fetus. Placental lactogens, such as Pl2, bind the prolactin receptor and elicit responses indistinguishable from those of prolactin (25, 32). Surges in prolactin secretion characterize the first half of pregnancy in rodents but cease with the onset of Pl2 production (21). Because serum Pl2 levels markedly exceed those of prolactin, Pl2 effectively replaces prolactin in pregnant mice from mid- to late gestation (21). Pl2, in turn, engages prolactin receptors on pancreatic β-cells to drive the adaptive increase in β-cell insulin secretion required to maintain maternal glucose homeostasis during pregnancy.

A defect in β-cell adaptation during pregnancy creates a permissive environment for the development of GDM (33, 34, 41, 42). Consistent with this, *Glut1^d/d^* females exhibit a striking onset of glucose intolerance during pregnancy, a defining feature of GDM. The reduced Pl2 levels in Glut1 mutants likely impair the adaptive expansion of β-cell insulin secretion, precipitating maternal hyperglycemia. A model describing this study is shown in Figure 13. Existing animal models of GDM have important limitations. Chemical models destroy pancreatic β-cells and produce a phenotype more consistent with type 1 diabetes than with GDM (35), and most genetic models target pancreatic genes, again recapitulating type 1 diabetes or non-pregnancy-specific glucose intolerance (33, 35). To our knowledge, the *Glut1^d/d^* model, in which an exclusively uterine perturbation triggers spontaneous maternal hyperglycemia restricted to pregnancy, represents one of the most clinically relevant genetic mouse models of GDM described to date, and it offers a tractable system for dissecting how endometrial metabolic dysfunction can give rise to maternal metabolic disease (41, 42).

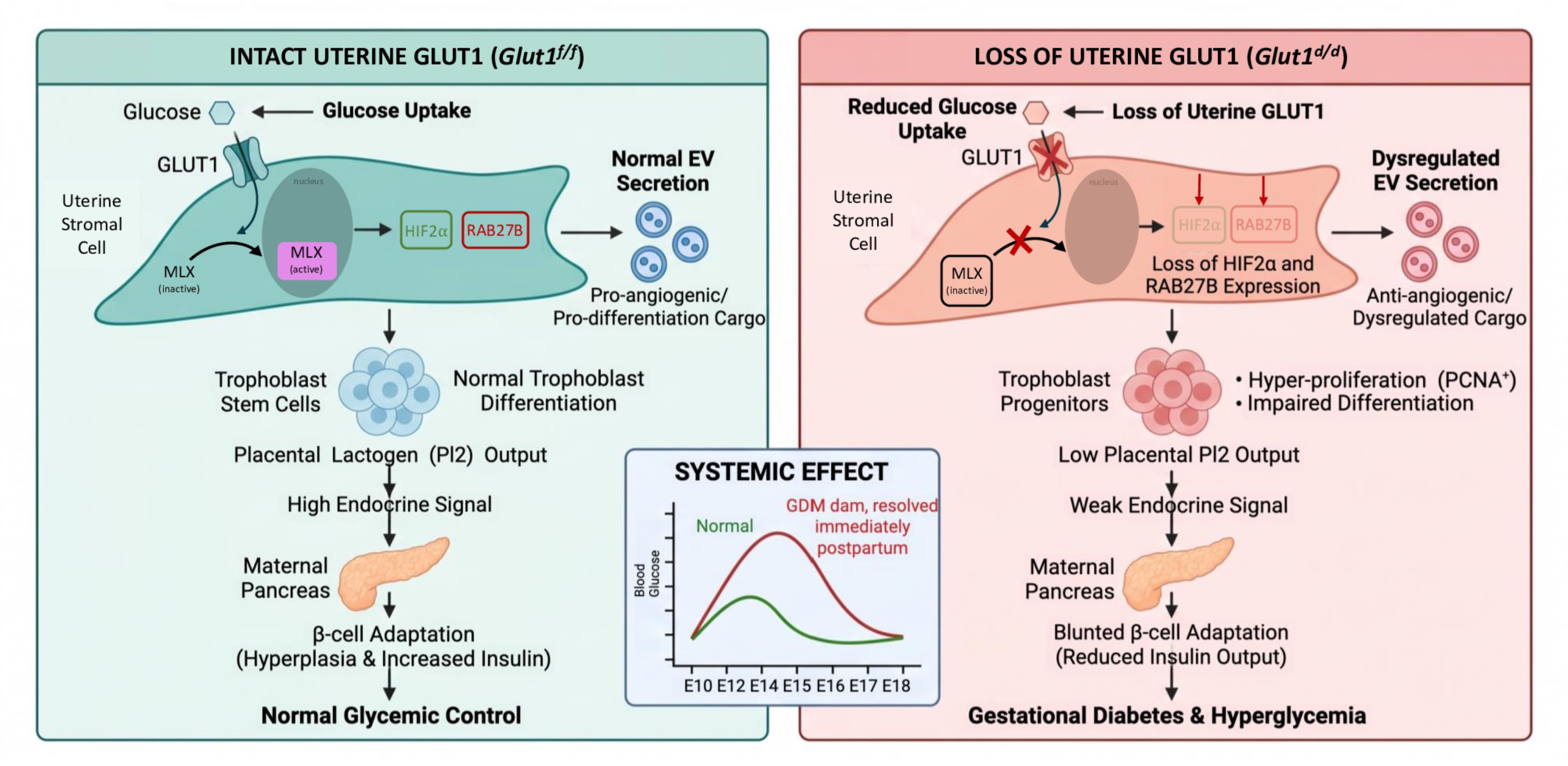

## MATERIALS AND METHODS

### Animals

Mice were maintained in the animal facility at the University of Illinois College of Veterinary Medicine in accordance with institutional guidelines for the care and use of laboratory animals and National Institutes of Health standards. Mice were housed at 22 °C on a 12L:12D cycle, with food and water available ad libitum. The Institutional Animal Use and Care Committee at the University of Illinois at Urbana–Champaign approved all procedures involving animal care, euthanasia, and tissue collection.

Conditional Glut1-null mice (*Glut1^d/d^*) were generated by crossing mice harboring a “floxed” Glut1 gene (*Glut1^f/f^*) with *Pgr-Cre* knock-in mice. *Glut1^f/f^* mice were obtained from the Jackson Laboratory. Pgr-Cre knock-in mice were provided by Drs. Francesco J. DeMayo and John P. Lydon (14). This strategy has been used extensively to ablate “floxed” genes in tissues expressing PGR (36, 37).

### Fertility assessments, timed pregnancies, and tissue collection

To test fertility, *Glut1^f/f^* and *Glut1^d/d^*females of reproductive age (7-8 weeks) were paired with fertile wild-type males for six months. The total number of pups per litter and the number of pregnancies during this period were recorded. For timed pregnancies, females were mated with adult wild-type males of known fertility. The identification of a copulatory plug was designated as day 1 of pregnancy. For tissue collection, animals were euthanized by CO₂ asphyxiation. Uteri were collected at the indicated time points and either fixed in 10% (vol/vol) neutral-buffered formalin (NBF) for histology or flash-frozen in liquid N₂ for RNA isolation or frozen sectioning.

### Primary mouse endometrial stromal cell isolation and induction of decidualization

Mouse endometrial stromal cells (MESC) were isolated from uteri on day 4 of pregnancy as previously described (30). Briefly, uteri were cut open and digested in 5 mL/uteri of 1× HBSS containing 6 g/L dispase and 25 g/L pancreatin for 45 min at room temperature, followed by 15 min at 37 °C, with occasional mixing. The supernatant containing epithelial cells was removed, and the remaining tissue fragments were washed with HBSS containing 10% (vol/vol) heat-inactivated FBS to stop the digestion. Tissue fragments were washed twice with 1× HBSS and then digested in 5 mL/uteri of 1× HBSS containing 0.5 g/L collagenase for 1 h at 37 °C. Digestion was stopped with 5 mL of HBSS containing 10% FBS. The tubes were vortexed for 10-15 s, and the supernatant was passed through an 80-µm gauze filter (Millipore) into a new tube to remove undigested myometrial fragments. Endometrial cells were pelleted at 430 × g for 5 min, washed with HBSS, and resuspended in DMEM/F12 supplemented with 2% FBS, 100 units/L penicillin, 0.1 g/L streptomycin, 1.25 mg/L Fungizone, 10 nM E2, and 1 µM P4. Live cells were assessed by trypan blue staining using a hemocytometer, and MESCs were seeded in 6-well plates at 5 × 10⁵ cells/well. Unattached cells were removed by HBSS washes, and culture was continued in fresh medium. At the end of the culture, cells were detached, counted using trypan blue and a hemocytometer, and stored at −80 °C for RNA extraction or immunocytochemical analysis.

### Quantitation of EVs by Microfluidic Resistive Pulse Sensing

MESCs were grown in DMEM/F-12 supplemented with 5% exosome-depleted FBS to harvest EVs. In vitro differentiation was stimulated with 1 µM progesterone and 10 nM 17β-estradiol (Sigma-Aldrich). Cell-conditioned medium was collected and centrifuged at 3000 g for 10 min to obtain a cell-free specimen. The supernatant was stored at −80 °C or directly extracted using the miRCURY Exosome Cell/Urine/CSF kit (Qiagen). EV pellets were resuspended in PBS and quantified by microfluidic resistive pulse sensing (MRPS) as described (18, 19). MRPS was performed using disposable microfluidic cartridges; this study used C-300 (50-300 nm) and C-400 (65-400 nm) cartridges. A 3–5 µL bolus of EV solution was injected into the cartridge reservoir. As particles flow through the MRPS aperture, they transiently alter the constriction’s electrical resistance by an amount proportional to the ratio of nanoparticle volume to aperture volume; therefore, EV concentration is accurately determined from these changes. MRPS reports particle concentration (p/mL) and particle diameter (nm). Data are presented as mean concentrations of repeated measurements on independent samples.

### Mass Spectrometry

Directly following isolation, extracellular vesicle (EV) pellets isolated from *Glut1^f/f^* and *Glut1^d/d^* stromal cells conditioned media were submitted to the Mass Spectrometry Laboratory at the University of Illinois at Urbana-Champaign for proteomic analysis. EVs were lysed in a lysis buffer comprising 6 M guanidinium chloride and 0.1% sodium deoxycholate in 100 mM triethylammonium bicarbonate. Aliquots were reserved for protein quantification via bicinchoninic acid (BCA) assay (Pierce; Thermo Fisher Scientific). The remaining sample volume was incubated at 95 °C for 15 min to ensure complete protein denaturation and facilitate the reduction and alkylation of disulfide bonds.

Proteins were sequentially digested using 500 ng of LysC (Wako) at 30°C for 3 h, followed by 500 ng of trypsin (Pierce) at 37°C overnight. The resulting peptides were acidified, desalted, and dried under vacuum centrifugation. Equal peptide masses across experimental groups were subsequently injected into an UltiMate 3000 RSLCnano system (Thermo Scientific) operating at a constant flow rate of 300 nL/min. Following peptide separation, standard column washing and regeneration protocols were executed.

Mass spectrometry analysis was conducted using data-dependent acquisition. Full MS1 scans were acquired across a mass-to-charge ratio (m/z) range of 350 to 1500 at a resolution of 120,000 (maximum injection time [IT]: 50 ms; automatic gain control [AGC] target: 3e6). The top 15 most abundant precursor ions from each MS1 scan were selected for higher-energy C-trap dissociation (HCD) fragmentation at a normalized collision energy (NCE) of 30. Subsequent MS2 scans were recorded at a resolution of 15,000 using a 1.2 m/z isolation window (maximum IT: 30 ms; AGC target: 5e4). To maximize proteome coverage and minimize redundant sampling of high-abundance ions, isotope exclusion was enabled, and the dynamic exclusion duration was set to 60 s.

Liquid chromatography-mass spectrometry (LC-MS) raw files were searched using Mascot v2.8.0 (Matrix Science) against the Mus musculus UniProt reference proteome (55,192 sequences). Search parameters specified tryptic digestion allowing a maximum of two missed cleavages, with the statistical significance threshold for protein identification set at P < 0.05. Label-free quantification (LFQ) was performed using Mascot Distiller v2.8.0 (Matrix Science) via an extracted ion chromatogram (XIC)-based average protocol, enforcing a strict minimum requirement of two unique peptides for protein abundance calculation.

### Immunohistochemistry (IHC)

Uterine tissues were processed for immunohistochemistry as previously described (9, 30). Briefly, paraffin-embedded tissues were sectioned at 5 µm and mounted. Sections were deparaffinized in xylene, rehydrated through ethanol, and rinsed in water. Antigen retrieval was performed in 0.1 M citrate buffer, pH 6.0, by microwave heating for 25 min. Endogenous peroxidase activity was quenched in 0.3% H₂O₂ in methanol for 15 min at room temperature. After washing in PBS for 15 min, sections were incubated in blocking solution for 1 h, then with primary antibodies overnight at 4 °C: anti-GLUT1, anti-PECAM1, anti-MLX, and anti-HIF2α. Placental sections were stained with anti-cytokeratin 8 (KRT8) and anti-placental lactogen 1 (PL1). Fluorescent secondary antibodies (rhodamine donkey anti-rabbit; 488 donkey anti-mouse; 488 donkey anti-goat) were purchased from Jackson ImmunoResearch. Fluoromount-G with DAPI was from eBiosciences. Images were acquired on an Olympus BX51 microscope with a Jenoptik ProgRes C14 digital camera (1.4 megapixel CCD) and processed and merged using Adobe Photoshop Extended CS6 (Adobe Systems).

### Quantitative real-time PCR (qPCR)

Total RNA was isolated from uteri, ovaries, and cells using a standard TRIzol-based protocol. RNA concentration was determined at 260 nm using a Nanodrop ND1000 spectrophotometer (Nanodrop Technologies). RNA was reverse transcribed using the High Capacity cDNA Reverse Transcription kit (Applied Biosystems) according to the manufacturer’s instructions. Gene-specific primers were used for real-time qPCR with SYBR-green master mix (Applied Biosystems) on a 7500 Applied Biosystems Real-time PCR instrument. The mean threshold cycle (Ct) was calculated from three replicates per sample. The normalized ΔCt in each sample was calculated as the mean Ct of the target gene minus the mean Ct of the reference gene, and ΔΔCt as the difference between control and mutant ΔCt values. Fold change was calculated using the 2^⁻ΔΔCt^ model (9, 30). Mean fold induction and SEM were calculated from at least three independent experiments. RPLP0 (36B4) was used as the reference gene. Data are reported as mean fold induction ± SEM.

### RNA sequencing analysis

Decidua devoid of embryo were isolated from *Glut1^f/f^*and *Glut1^d/d^* mice on day 7 of pregnancy. Total RNA was extracted using a standard TRIzol-based protocol (14). RNA integrity was verified using an Agilent 2100 Bioanalyzer (Agilent Technologies, Santa Clara, CA, USA), and sequencing was performed at the Biotechnology Center of the University of Illinois, Urbana–Champaign. Genes with a relative fold change >2 were sorted by gene ontology and pathway analysis.

### Blood glucose measurement

Blood glucose levels were measured using a FreeStyle Lite blood glucose monitoring system and the corresponding test strips. Each mouse was fasted for 12-16 h prior to measurement. Blood was collected by pricking the tail with a needle. Measurements are reported in milligrams per deciliter (mg/dL).

### Measurement of Serum Insulin Levels

Circulating insulin concentrations were quantified in blood serum collected from *Glut^f/f^* and *Glut^d/d^* mice at gestational day 18 using a commercial Mouse Insulin Sandwich ELISA Kit (Cat. No. KE10089; Proteintech Group, Inc.) according to the manufacturer’s instructions. The analytical sensitivity of the assay was 1.3 pg/mL, with a dynamic detection range of 6.25 to 400 pg/mL.

Prior to the assay, serum samples were thawed and diluted 1:64 or 1:128 in Sample Diluent PT 4 to ensure that raw optical density (OD) readings fell within the linear range of the standard curve. Recombinant mouse insulin protein standards were prepared via a two-fold serial dilution to generate a 7-point standard curve ranging from 400 pg/mL down to 6.25 pg/mL; Sample Diluent served as the zero standard (0 pg/mL).

The assay was performed at room temperature using pre-coated 96-well microplates. Standards and diluted serum samples 100 μL per well) were added in duplicate to the plate and incubated for 2 h at 37 °C. Following incubation, the liquid was aspirated, and the wells were washed four times with Wash Buffer (350-400 μL per well). Next, 100 μL of a 1x Detection Antibody solution was added to each well, and the plate was incubated for 1 h at 37°C, followed by a second four-fold wash cycle. Horseradish peroxidase (HRP)-conjugated secondary antibody (1x, 100 μL per well) was then introduced and incubated for 40 min at 37 °C.

After a final four-fold wash step to remove unbound conjugate, signal development was initiated by adding 100 μL of tetramethylbenzidine (TMB) substrate solution. The plate was incubated in the dark at 37 °C for 15 to 20 min, turning the solution blue in proportion to the bound insulin concentration. Enzymatic color development was subsequently quenched by adding 100 μL of sulfuric acid-containing Stop Solution per well, shifting the color to yellow.

Absorbance was measured immediately within 5 min on a microplate reader at 450 nm, with wavelength correction set at 630 nm to account for optical imperfections in the plate. Absolute insulin concentrations were calculated by subtracting the average zero-standard blank OD from all readings and fitting the corrected data to a four-parameter logistic (4-PL) curve-fit regression model. Final values were adjusted by multiplying the calculated concentrations by the respective sample dilution factors.

### Statistical analyses

Statistical analyses were performed as previously described (30). Experimental data were collected from at least four independent samples under identical conditions. Numerical data are expressed as mean ± SEM. Statistical analysis used a two-tailed Student’s t-test and a one-way ANOVA, followed by Tukey’s post hoc test for multiple comparisons. Variance equality was assessed for all numerical data to determine whether parametric or non-parametric tests were appropriate. Data were considered statistically significant at p ≤ 0.05 and are indicated by asterisks in the figures. All data were analyzed and plotted using GraphPad Prism 9.0 (GraphPad Software).

